# Fitness costs associated with spinetoram resistance in *Spodoptera frugiperda* is driven by host plants

**DOI:** 10.1101/2021.11.26.470136

**Authors:** Rubens H. Kanno, Aline S. Guidolin, Fernando E. O. Padovez, Juliana G. Rodrigues, Celso Omoto

**Author notes:** Corresponding to Rubens H. Kanno.

## Abstract

Insecticide resistance is usually associated with fitness costs. The magnitude of fitness costs is affected by environmental and ecological factors. Here, we explored how host plants could affect fitness costs associated with insecticide resistance. Initially, spinetoram-resistant (RR) and susceptible (SS) strains of *Spodoptera frugiperda* were selected using F_2_ screen from a population collected in São Desidério, Bahia State, Brazil in 2018. Besides de RR and SS strains, fitness costs were also assessed for a heterozygous strain (RS). Life-history traits were evaluated to estimate population growth parameters of neonate larvae of each strain fed on corn, soybean and cotton plants. Compared to the SS strain, the relative fitness of the RR strain, based on intrinsic rate of population increase, was 1.06, 0.84 and 0.67 on plants of corn, soybean and cotton respectively. The relative fitness of the RS strain was similar to the SS strain regardless the host plant, suggesting a recessive fitness cost. No differences were found between the strains fed on corn plants. The larval development time was greater for RR strain fed on soybean and cotton plants compared to RS and SS strain. Low survival rate and fecundity of the RR strain were found when larvae fed on plants of soybean and cotton. The results of this study demonstrated that fitness costs of spinetoram resistance in *S. frugiperda* depend strongly on the host plants that *S. frugiperda* larvae fed on. Such information can be used to design resistance management strategies considering the host plants of the agricultural landscape.

**Key messages:**

- The presence of fitness costs associated with resistance can be exploited in resistance management strategies.
- Host plant influences the fitness costs associated with spinetoram resistance in *S. frugiperda*.
- Information considering the host plants in an agricultural landscape is essential to design effective resistance management programs.

## Introduction

The fall armyworm, *Spodoptera frugiperda* (Lepidoptera: Noctuidae), is recognized as one of the most important agricultural damaging pests. Recently, the notoriety of this pest has increased because it has been reported as an invasive pest in many countries of the African, Asian and Oceanian continent (Goergen et al. 2016; Baloch et al. 2020; CABI 2021). *Spodoptera frugiperda* is a highly polyphagous pest with a wide range of host plant species (Montezano et al. 2018). The ability of this pest to explore different host plant species is one of the major factors of its success to colonize new areas. The control of *S. frugiperda* relies mainly on the use of chemical insecticides, but the extensive use of this control tactics resulted in resistance cases to many groups of insecticides (Diez-Rodríguez and Omoto 2001; Carvalho et al. 2013; Nascimento et al. 2016; Okuma et al. 2018; Bolzan et al. 2019; Lira et al. 2020; Muraro et al. 2021; Garlet et al. 2021a), challenging the management of this pest.

The spinosyns are a broad-spectrum insecticides derived from natural fermentation of *Saccharopolyspora spinosa* (Sparks et al. 2001). This group of insecticide represented by two chemical molecules, spinosad and spinetoram, acts as allosteric modulators of nicotinic acetylcholine receptor (Crouse et al. 2001; Salgado et al. 2010; Dripps et al. 2011). These insecticides have demonstrated high efficiency against many insect pests (Salgado et al. 2010; Dripps et al. 2011). However, resistance has already been documented to spinosyns in some insect pests (Sparks et al. 2012; Mota-Sanchez and Wise 2021), including *S. frugiperda* with resistance ratio >890-fold in Brazil (Okuma et al. 2018; Lira et al. 2020). The high levels of resistance for spinosyns in *S. frugiperda* indicates the elevated risk of resistance evolution in this pest and the urgency in implementing insect resistance management strategies.

One of the main insect resistance management strategies is based on the assumption of fitness costs associated with resistance (Roush and McKenzie 1987; Gassmann et al. 2009; Kliot and Ghanim 2012). The fitness costs of insecticide resistance can be understood as a significant disadvantage of resistant individuals compared with its susceptible counterpart in the absence of insecticides (Kliot; Ghanim, 2012). In general, a fitness costs are associated with spinosyn resistance. A significant reduction in the rate of survival to adulthood and a lower reproductive rate were found in a spinosad resistant strain of *S. frugiperda* (Okuma et al. 2018). Fitness cost of spinosad resistance has also been reported for other pests such as *Chloridea virescens* (Wyss et al. 2003), *Plutella xylostella* (Li et al. 2007b), *Helicoverpa armigera* (Wang et al. 2010), *Spodoptera litura* (Rehan and Freed 2015), *Phenacoccus solenopsis* (Afzal and Shad 2017) and *Ceratitis capitata* (Guillem-Amat et al. 2020). Spinetoram is a more recent insecticide molecule in the market than spinosad, and so far only one study about the fitness costs associated to spinetoram resistance was reported in *Thrips hawaiiensis* (Fu et al. 2018), therefore such information remains unknown for *S. frugiperda*.

The magnitude of fitness costs associated with insecticide resistance can be influenced by various environmental and ecological factors (Gassmann et al. 2009). Different host plant species and allelochemicals could play an important role on fitness costs (Carrière et al. 2004; Janmaat and Myers 2005; Bird and Akhurst 2007; Raymond et al. 2007, 2011; Wang et al. 2016; Chen et al. 2018). The quality of the host plant can affect several insect development processes (Awmack and Leather 2002) and this is important for population increase and outbreaks of insect pests, especially those that can feed on a large range of host plants (Kennedy and Storer 2000; Sivakoff et al. 2013). Therefore, the understanding of the interaction between the host plant and fitness costs associated with insecticide resistance is essential to implement effective resistance management strategies.

Previous works about the interaction of host plants and fitness costs have focused on Bt resistance (Janmaat and Myers 2005; Bird and Akhurst 2007; Raymond et al. 2007, 2011; Wang et al. 2016; Chen et al. 2018). To date there is just one study conducted by Garlet et al. (2021b) exploring this interaction with chemical insecticide resistance. The recent documentation of spinosyn resistance in *S. frugiperda* (Okuma et al. 2018; Lira et al. 2020) and the broad host range of this pest provides us the unique opportunity to investigate the effect of different host plants on fitness costs associated with spinosyn resistance. Corn, soybean and cotton are three of the most economically important crops in Brazil (Buainain et al. 2019), and its intensive cultivation offers an ideal scenario for the development of *S. frugiperda* populations across the year, since this pest feed on all these crops (Barros et al. 2010). In this context, we aimed to assess the fitness costs of spinetoram resistance in *S. frugiperda* by comparing several biological parameters and constructing fertility life-tables of the resistant, susceptible and heterozygous strains fed on plants of corn, soybean and cotton. The findings of this study will contribute to improve insecticide resistance management strategies of *S. frugiperda* and understanding how host plants can affect the population growth under different agricultural landscapes.

## Material and methods

### Insect strains

The spinosyn resistant strain (RR) of *S. frugiperda* was selected from a field population collected in São Desidério – BA in 2018. The F_2_ screen method was used to obtain the resistant strain (Andow and Alstad 1998). To investigate the fitness cost of spinosyn resistance in strains with same genetic background, a spinosyn susceptible strain (SS) was obtained from the same field population that originated the resistant strain. The selection for susceptibility was conducted establishing pair-mated adults from the field population. The larvae from the F_1_ progeny of each couple were separated in two groups, one group was tested at the diagnostic concentration of 5.6 μg a.i. spinetoram/ml (sufficient to kill the susceptible individuals) (Lira et al. 2020) and another group remained to establish the susceptible strain if the respective progeny had a 100% mortality. Both strains was reared on artificial diet (Kasten Jr et al. 1978). The heterozygous strain (RS) was obtained from the cross of RR females and SS males. Only one heterozygous strain was established because the inheritance pattern of spinosyn resistance in *S. frugiperda* is autosomal and both heterozygotes obtained from reciprocal crosses showed a similar mortality to spinosad and spinetoram (Okuma et al. 2018; Lira et al. 2020).

Concentration-response curves were performed to characterize the susceptibility of the SS and RR strains. In addition, a laboratory susceptible strain (SS-Lab), which is maintained in laboratory for more than 25 years without the selection pressure by any insecticide or Bt proteins, was used to validate the SS strain as a susceptible strain. Diet overlay bioassays were conducted in 24-well acrylic plates containing 1.25 ml of artificial diet in each well (1.9 cm^2^ area). All strains were tested in eight logarithmically spaced concentrations, ranging from 0.1 to 5.6 μg a.i. spinetoram/ml for SS and SS-lab strains and from 180 to 5,600 μg a.i. spinetoram/ml for the RR strain. The different concentrations of spinetoram were obtained by the dilution of the formulated insecticide (Exalt^®^ 120g a.i./l) in distilled water with the addition of 0.1% (v/v) of the surfactant Triton X-100 (Sigma Aldrich Brasil Ltda). In each well, 30 μl of the insecticide solution was applied. One third instar larvae were infested in each well after the drying of the insecticide solution. The mortality was assessed after 48h and larvae that did not show coordinated movement when prodded were considered dead.

### Fitness cost assessment bioassays

Fitness costs associated with spinosyn resistance in *S. frugiperda* were investigated on plants of corn, soybean and cotton. Plants of non-Bt hybrid corn (3700 RR2), non-Bt soybean (95R51) and non-Bt cotton (FM 944GL) were cultivated in 12l pots in a greenhouse. The SS and RR strains were reared for one generation in each host plant before the fitness cost bioassays to eliminate possible effects of changing the food source.

The bioassays were performed with leaves of corn from V4 to V8 growth stages, leaves of soybean from V3 to V6 and leaves of cotton from squaring phenological stage. The leaves of each plant were cut into pieces (approximately 4 cm^2^) and placed over a gelled mixture of 2.5% agar-water in 16-well plastic trays (Advento do Brasil). One neonate (<24h old) from each strain was infested in each well and reared until the 6^th^ instar. Then, the larvae were transferred to another 16-well plastic tray containing vermiculite and leaves of the respective host plants for development until the pupal stage. The leaves were changed every day. For each strain, 160 larvae were tested (10 replicates of 16 larvae) per host plant. The following parameters was evaluated: duration and survival rates of the egg, larval, pupal and egg-adult period; pupal weight; sex ratio; male and female longevity; duration of the period of preoviposition, oviposition and postoviposition, fecundity and fertility. To determine the male and female longevity, duration of the period of preoviposition, oviposition and postoviposition, fecundity and fertility, 20 couples per strain were kept in PVC cages internally coated with paper for mating and oviposition. The adults were fed with 10% honey solution soaked in cotton. The embryonic period and survival were evaluated in 100 eggs of the second oviposition of each pair. All parameters were evaluated in daily observations. The bioassay trays and the insects were maintained in rearing rooms under controlled conditions of 25 ± 2 °C, 70% relative humidity and a photoperiod of 14:10 (L:D) h.

### Statistical analysis

The mortality data of concentration-response curves were fitted to a generalized linear model with binomial distribution and probit as function link. The LC_50_s and the respective confidence intervals were estimated using the function *dose*.*p* from MASS package (Venables and Ripley 2002).

The data of fitness cost bioassays were analyzed using a generalized linear model (glm) according to the distribution of each data. The pupal weight and fertility life table parameters data were fitted to a glm for Gaussian distribution; data of the development time of egg, larvae, pupae and egg-adult period and the number of eggs were fitted to a glm for quasi-Poisson distribution; survival rates of egg, larvae, pupae and egg-adult period were fitted to a glm for quasibinomial distribution. The goodness of fit was verified using the *hnp* package (Moral et al. 2017). The fertility life table, which includes the net reproductive rate (*R*_*0*_), the mean length of a generation (*T*), the intrinsic rate of population increase (*rm*) and the finite rate of population increase (*λ*), was estimated using the *lifetable*.*r* procedure (Maia et al. 2014). The relative fitness was calculated by dividing the *rm* values of the RR or RS strain by the *rm* value of the SS strain. ANOVA was performed to verify the effect of each factor (strain; host plant) and their interaction using glm for each evaluated parameter, followed by a multiple pairwise comparison (Tukey test) using the *lsmeans* package. All statistical analysis was performed in R Software (R Core Team, 2020).

## Results

### Susceptibility of *Spodoptera frugiperda* strains to spinetoram

The SS and SS-Lab strains had a similar susceptibility to spinetoram (Table 1). The LC_50_ value of SS strain was 1.0 μg ml^-1^, while the SS-Lab strain presented a LC_50_ of 0.8 μg ml^-1^, indicating a low variation of 1.2-fold. The LC_50_ value of RR strain was 776.9 μg ml^-1^, which results in a resistance ratio of 951.1-fold when compared to the SS-Lab strain and a resistance ratio of 776.9-fold when compared to the SS strain.

**Table 1.** Susceptibility of *Spodoptera frugiperda* strains to spinetoram

| Strain | n<sup>a</sup> | Slope (± SE) | $LC_{50}$ (CI 95%)<sup>b</sup> | $\chi^2$ | df<sup>c</sup> | RR<sup>d</sup> |
| --- | --- | --- | --- | --- | --- | --- |
| SS-Lab | 648 | $2.4 \pm 0.2$ | $0.8 (0.7 - 0.9)$ | 9.1 | 5 | - |
| SS | 598 | $2.2 \pm 0.2$ | $1.0 (0.8 - 1.2)$ | 8.1 | 5 | 1.2 |
| RR | 693 | $2.6 \pm 0.2$ | $776.9 (685.7 - 880.3)$ | 9.8 | 5 | 951.1 |

<sup>a</sup> number of larvae tested; <sup>b</sup> lethal concentration ( $\mu\text{g ml}^{-1}$ ) of applied insecticide solution that kills 50% of the individuals; <sup>c</sup> degrees of freedom; <sup>d</sup> Resistance ratio: $LC_{50}$ of the tested strain/ $LC_{50}$ of the susceptible reference strain.

### Survivorship of *Spodoptera frugiperda* strains on plants of corn, soybean and cotton

The effect of strain, host plant and their interaction on survivorship of egg stage were not significant (F = 2.64, df = 2, 87, p = 0.07 for strain; F = 2.76, df = 2, 85, p = 0.06 for host plant; F = 2.11, df = 4, 81, p = 0.08 for interaction). A high survivorship of egg stage (>92%) was observed in all strains regardless the host plant (Fig. 1).

**Fig. 1.**
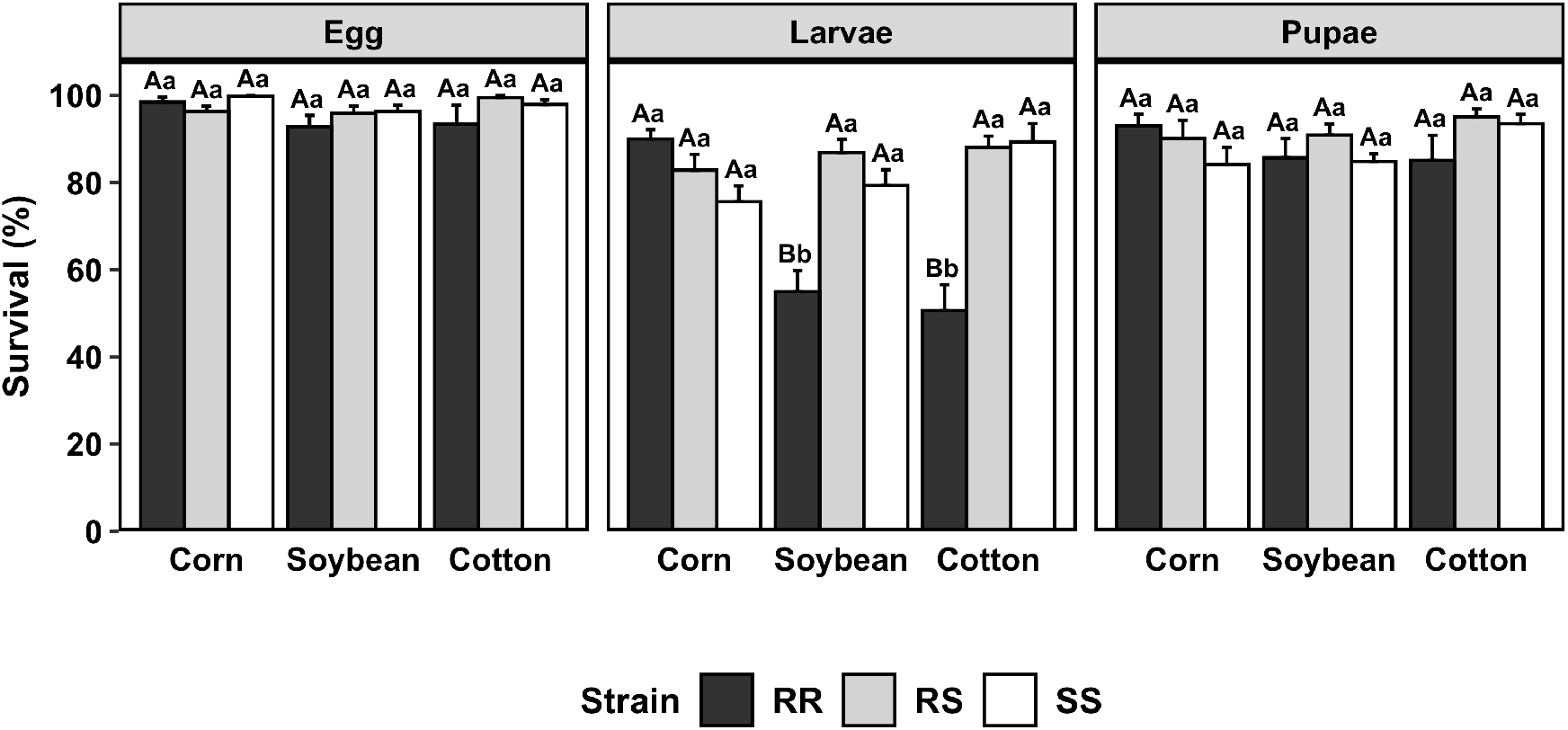
Survival rate of different life stages of *Spodoptera frugiperda* strains on plants of corn, soybean and cotton. Bar height represents the means of each treatment and error bars represents the standard error of the mean. Different lowercase letters indicate a significant difference between *S. frugiperda* strains in each host plant and uppercase letters indicates significant difference for the same strain on different host plants (Tukey test, p<0.05).

The larval stage was affected by the strain, host plant and their interaction (F = 21.20, df = 2, 87, p < 0.05 for strain; F = 4.44, df = 2, 85, p < 0.05 for host plant; F = 12.24, df = 4, 81, p < 0.05 for interaction). The SS and RS strains presented a high survivorship (>75%) on the three host plants and did not differ between them. The difference was observed on the survivorship of the larval stage of the RR strain in the different host plants. In corn plants, the RR strain had a larval survivorship of 90%, while on soybean and cotton plants the survivorship was 55 and 50.62%, respectively (Fig. 1).

No difference was observed on survivorship of the pupal stage among the SS, RS and RR strains and between the three host plants evaluated (F = 2.10, df = 2, 87, p = 0.12 for strain; F = 1.19, df = 2, 85, p = 0.30 for host plant; F = 1.96, df = 4, 81, p = 0.10 for interaction) (Fig. 1).

### Development time of *Spodoptera frugiperda* strains on plants of corn, soybean and cotton

The effect of strain and the interaction between strain and host plant were significant on the duration of the egg stage (F = 8.60, df = 2, 87, p < 0.05 for strain; F = 10.20, df = 4, 81, p <0.05 for interaction). The effect of host plant was not significant (F = 2.04, df = 2, 85, p = 0.13). The duration of the egg stage ranged from 3 to 4 days in all strains evaluated. Difference between the three strains was observed only in soybean and cotton plants (Fig. 2).

**Fig. 2.**
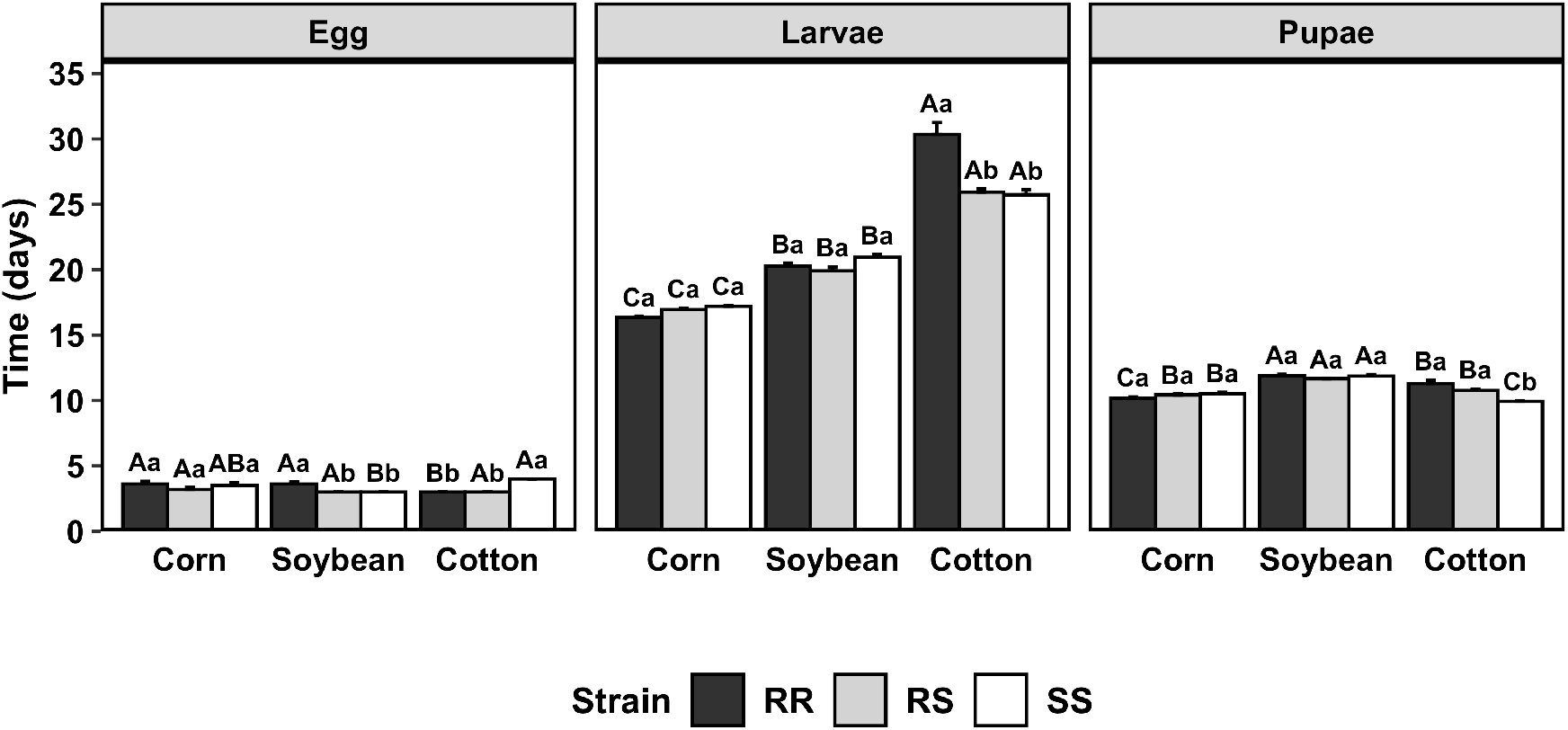
Development time of different life stages of *Spodoptera frugiperda* strains on plants of corn, soybean and cotton. Bar height represents the means of each treatment and error bars represents the standard error of the mean. Different lowercase letters indicate a significant difference between *S. frugiperda* strains in each host plant and uppercase letters indicates significant difference for the same strain on different host plants (Tukey test, p<0.05).

The duration of larval stage was affected by the strain, host plant and their interaction (F = 8.93, df = 2, 87, p < 0.05 for strain; F = 948.44, df = 2, 85, p < 0.05 for host plant; F = 22.19, df = 4, 81, p < 0.05 for interaction). The longest duration of the larval stage was observed when the strains were reared on cotton plants. On cotton, the RR strain presented a duration of 30.36 days, differing from SS and RS strains which had a duration of 25.74 and 25.93 days, respectively. No difference was observed among the strains when they were reared on corn and soybean plants. The larval stage was 3.55 – 3.74 days longer on soybean plants compared to corn plants (Fig. 2).

The strain, host plant and their interaction had a significant effect on the duration of pupal stage (F = 5.72, df = 2, 87, p < 0.05 for strain; F = 107.86, df = 2, 85, p < 0.05 for host plant; F = 13.72, df = 4, 81, p < 0.05 for interaction). Difference on duration of pupal stage among the strains was observed only on cotton plants, which the SS strain had the shorter duration (9.93 days) differing from RS and RR with had a duration of 10.76 and 11.31 days, respectively. The longer duration of the strains between the plants was observed on soybean, with a duration ranging from 11.67 to 11.88 days among the strains (Fig. 2).

### Pupal weight and fecundity of *Spodoptera frugiperda* strains on plants of corn, soybean and cotton

The effects of strain, host plant and their interaction were significant on pupal weight (F = 14.05, df = 2, 87, p < 0.05 for strain; F = 52.57, df = 2, 85, p < 0.05 for host plant; F = 34.41, df = 4, 81, p < 0.05 for interaction). The difference on pupal weight among the strains was observed on soybean and cotton plants. On soybean plants, the SS strain had a pupal weight of 188.23 mg, differing from RS and RR strains which had a pupal weight of 202.96 and 207.36 mg, respectively. On cotton plants, the RR strain had the lower pupal weight (158.77 mg) differing from SS (197.82 mg) and RS (193.82 mg) strains (Fig. 3A).

**Fig. 3.**
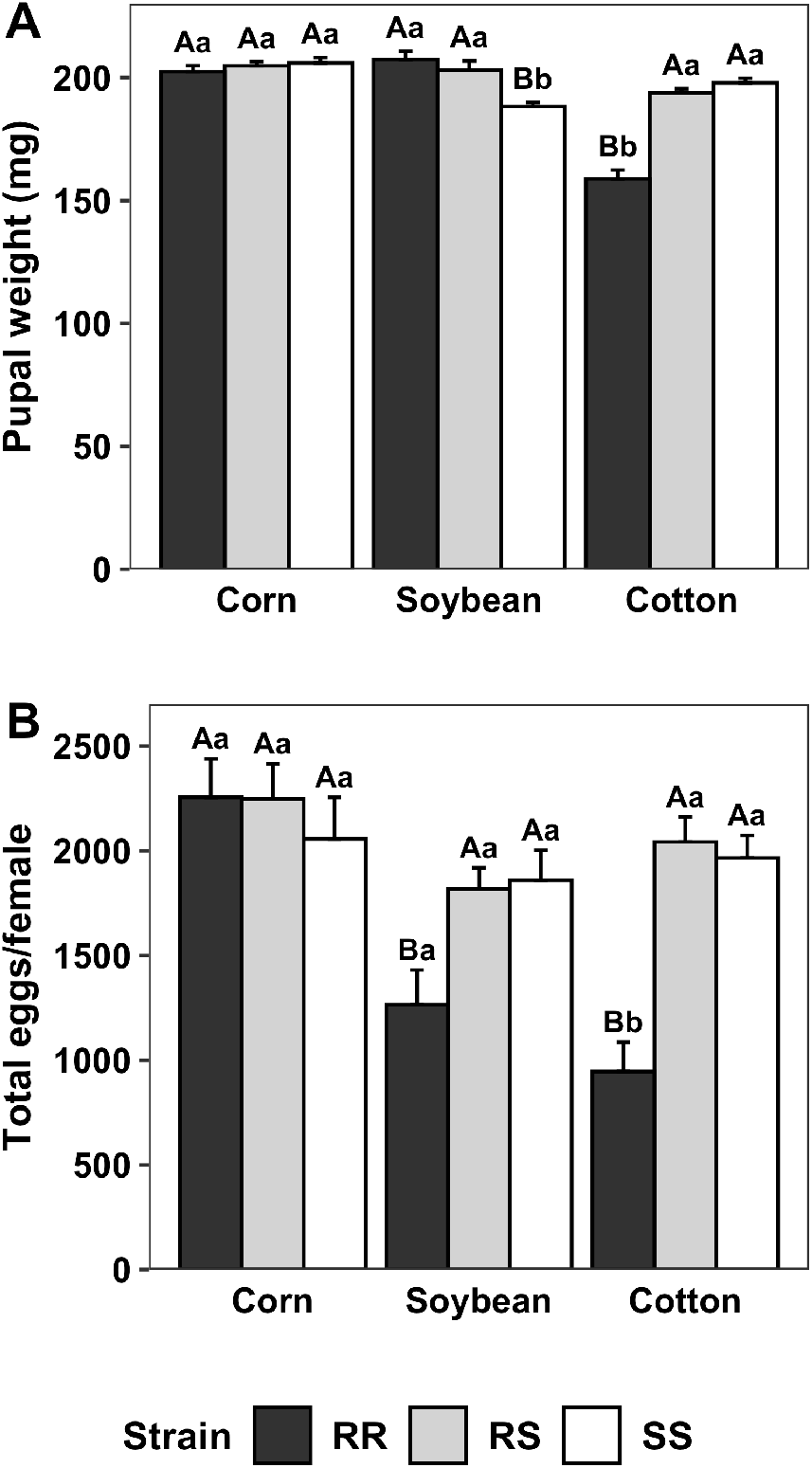
Biological parameters of *Spodoptera frugiperda* strains reared on plants of corn, soybean and cotton: (A) Pupal weight and (B) Number of total eggs per female. Bar height represents the means of each treatment and error bars represents the standard error of the mean. Different lowercase letters indicate a significant difference between *S. frugiperda* strains in each host plant and uppercase letters indicate significant difference for the same strain on different host plants (Tukey test, p<0.05).

The fecundity of *S. frugiperda* was affected by strain, host plant and their interaction (F = 9.38, df = 2, 152, p < 0.05 for strain; F = 11.63, df = 2, 150, p < 0.05 for host plant; F = 6.65, df = 4, 146, p < 0.05 for interaction). The number of total eggs laid per female did not differ among SS, RS and SS strains on corn plants. However, a large reduction was observed in the number of eggs laid by RR strain when it was reared on soybean and cotton plants. The SS and RS strain reared on soybean and cotton had a similar fecundity with no significant difference between them (Fig. 3B).

### Population growth parameters of *Spodoptera frugiperda* strains on plants of corn, soybean and cotton

Differences on population growth parameters were observed when the SS, RS and RR strains were reared on plants of corn, soybean and cotton (Table 2). The net reproductive rate (*R*_*0*_) was affected by strain, host plant and their interaction (F = 43.36, df = 2, 154, p < 0.05 for strain; F = 33.46, df = 2, 152, p < 0.05 for host plant; F = 20.04, df = 4, 148, p < 0.05 for interaction). The *R*_*0*_ values of the SS, RS and RR strains reared on corn ranged from 699.74 to 836.98, with no significant differences among them. A difference among strains was observed on plants of soybean and cotton. When reared on plants of soybean, the RR strain presented a *R*_*0*_ value of 239.33 while the *R*_*0*_ values of SS and RS strains were 580.02 and 637.84, respectively. In plants of cotton, the RR strain showed a *R*_*0*_ value of 144.90, differing from SS and RS strain which showed a *R*_*0*_ values of 786.30 and 813.85, respectively.

**Table 2.**
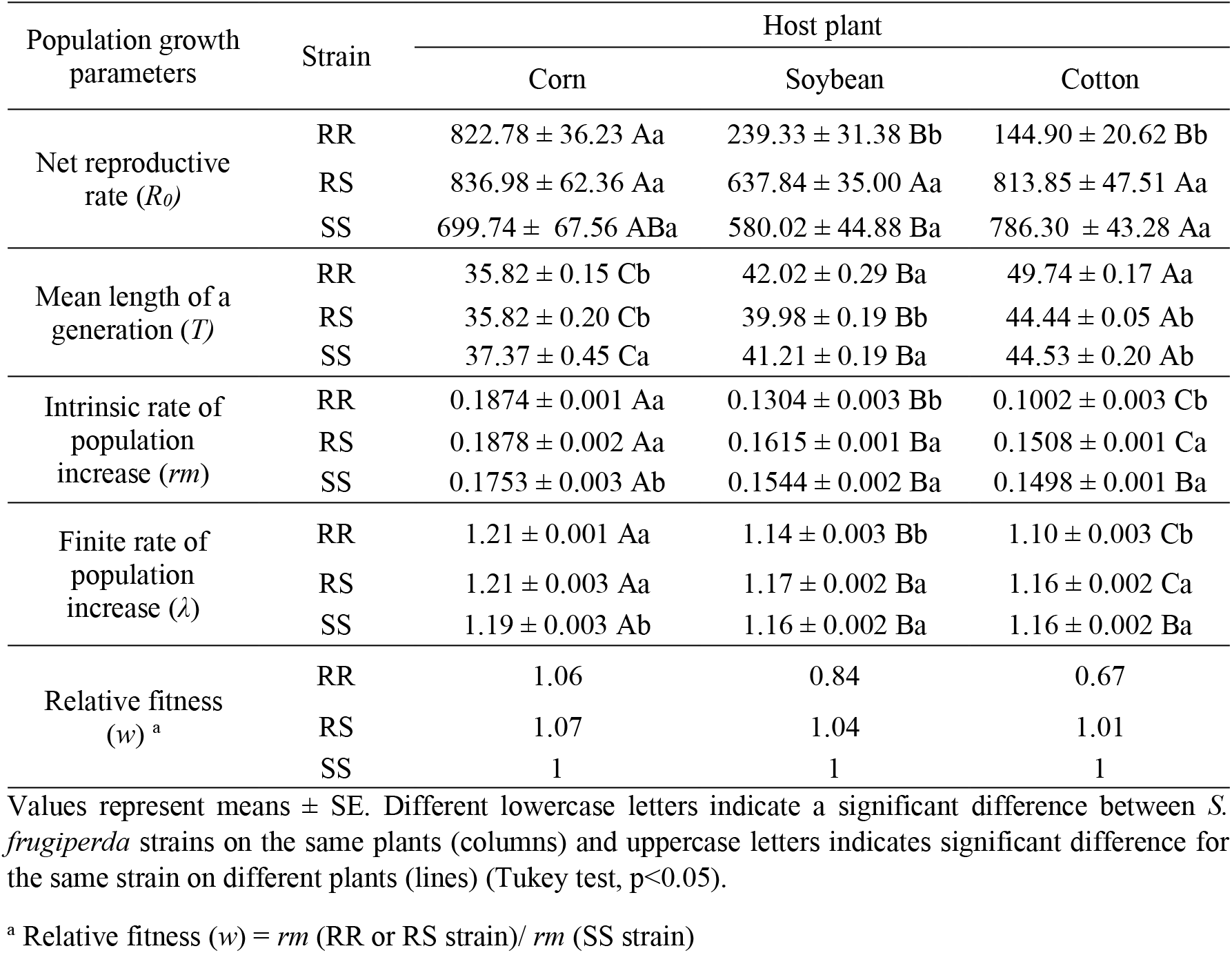
Population growth parameters of *S. frugiperda* strains reared on plants of corn, soybean and cotton

The mean length of a generation (*T*) was affected by strain, host plant and their interaction (F = 49.46, df = 2, 154, p < 0.05 for strain; F = 1431.17, df = 2, 152, p < 0.05 for host plant; F = 61.59, df = 4, 148, p < 0.05 for interaction). In corn plants, the duration of a generation was similar for RR and RS strains (35.82 days) and it was significantly shorter when compared to the SS strain (37.37 days). No differences were verified in the time of a generation between RR (42.02 days) and SS (41.01 days) strains on soybean plants, however these two strains differed from the RS strain (39.98 days). The longest durations of a generation of SS, RS and RR strains were observed for cotton plants. The mean length of a generation of SS and RS strains in cotton plants were 44.53 and 44.44 days, respectively, differing from the RR strain which presented a duration of 49.74 days.

The effects of strain, host plant and their interaction were all significant for the intrinsic rate of population increase (*rm*) (F = 98.84, df = 2, 154, p < 0.05 for strain; F = 448.49, df = 2, 152, p < 0.05 for host plant; F = 61.05, df = 4, 148, p < 0.05 for interaction) and for the finite rate of population increase (F = 94.48, df = 2, 154, p < 0.05 for strain; F = 452.26, df = 2, 152, p < 0.05 for host plant; F = 58.95, df = 4, 148, p < 0.05 for interaction). On corn plants, the SS strain presented the lowest intrinsic rate of population increase (0.1753) differing from RR (0.1874) and RS (0.1878) strains. While on soybean and cotton plants, the lowest *rm* was observed in the RR strain differing from the RS and SS strains. Similar pattern was verified for the finite rate of population increase (*λ*). The lowest finite rate of population increase was observed for the SS strain on corn, whereas on plants of soybean and cotton it was observed for the RR strain.

Considering the relative fitness (*w*) of the SS strain as a reference (*w* = 1), the relative fitness for the RR and RS strains on corn plants are 1.06 and 1.07, respectively. Decrease on the relative fitness were observed for the RR strain on plants of soybean (*w* = 0.84) and cotton (*w* = 0.67). Although, the RS strain did not show any reduction on the relative fitness when reared on soybean and cotton plants (*w* > 1 for both host plants). According to these results, the RR strain did not show a competitive disadvantage compared with the SS strain on plants of corn. However, a reduction of the competitiveness of the RR strain was observed on soybean and cotton plants when compared to the SS strain. The RS strain did not demonstrate any disadvantage compared to the SS strain on the three host plants evaluated, indicating that the fitness costs of spinetoram resistance in *S. frugiperda* are recessive independently of the host plant.

## Discussion

In this study, we demonstrated the effects of host plants on fitness cost associated with spinetoram resistance in *S. frugiperda* through the comparison of different biological parameters using strains with same genetic backgrounds. Thus, varying only the food source, no fitness costs associated to spinetoram were found in *S. frugiperda* reared on corn plants, but a significant fitness cost was observed in *S. frugiperda* resistant to spinetoram reared on soybean and cotton plants.

Recently, the importance of the genetic background have been highlighted on studies of fitness components (ffrench-Constant and Bass 2017; Lenormand et al. 2018; Freeman et al. 2021). The use of strains with same genetic backgrounds are particularly important in fitness cost studies because the impact of genetic variability of each strain could influence biological factors that are not associated with resistance (Kliot and Ghanim 2012; ffrench-Constant and Bass 2017). In this way, the use of near isogenic strains minimizes the overestimation of the presence of fitness cost associated with resistance and increases the probability to obtain information that reflect what is occurring in the field. However, the establishment of a near isogenic strain is time consuming and an alternative is to select a susceptible strain from the same field collection of the resistant strain (Freeman et al. 2021).

If the susceptible and the resistant strain are selected at the same time, both strains will suffer the laboratory rearing pressures from the same amount of time. Usually, laboratory susceptible strains, used as susceptibility references in several resistance studies, are commonly isolated from field populations and maintained under laboratory conditions for a long time. These strains are conditioned to artificial environments and present a reduced genetic variability, even though it preserves the susceptibility characteristics to pesticides which is a crucial factor in studies of resistance characterization, it also differs significantly from the original field population or from the resistant strain. The resistant (RR) and the susceptible (SS) strains of this work were selected from the same field population. Through concentration-response curves we validated this susceptible strain once it presented a similar susceptibility to the laboratory susceptible strain (SS-Lab).

The concentration-response curves also estimated a high resistance ratio for the resistant strain of *S. frugiperda* to spinetoram used in this study. Similar high resistance ratio has also been reported for spinosyn resistance by Okuma et al. (2018) and Lira et al. (2020). The increase in the number of highly resistant strains selected in a short period of time raises the hypothesis that the frequency of spinosyn resistance alleles in *S. frugiperda* populations in the field is rising, evidencing the need of the implementation of insect resistance management programs.

Fitness cost is one of the concepts that uphold an enduring resistance management program, our results showed that all strains (resistant, susceptible, and heterozygous) survived and completed their life cycles on plants of corn, soybean, and cotton. However, different patterns of development were observed among the three host plants evaluated. All strains developed fast and had a higher fecundity on corn compared to soybean and cotton. Therefore, the resistant management program establish for this species should be different regarding the host plant since the resistance development to spinetoram in corn crop tends to occur faster than soybean and cotton crops.

No significant fitness costs associated with spinetoram resistance on corn plants means that removing the selective pressure from the environment would not result in a decrease of spinetoram resistance allele frequency on corn plants, since there are no competitive disadvantage between resistant and susceptible individuals. On the other hand, a negative impact was verified on the life history traits of the RR strain on soybean and cotton plants, which had a significant reduction in the survival rate and the reproductive rate compared to the other strains. This information could be exploited to design effective resistance management strategies, for example a seasonal removal of spinosyn insecticides in the control of *S. frugiperda* and an adoption of insecticides with different modes of action in a rotation scheme, or other control tactic would aid to maintain the resistance of spinetoram at low frequencies.

The lack of fitness cost on corn plants could be related to the adaptation of *S. frugiperda* in this host plant. Silva et al. (2017) showed that corn and other grasses plants are the most suitable host plants for *S. frugiperda*. Although the preference for plants of the Poaceae family, this pest can complete its life cycle in any available plant in the absence of the preferred hosts (Barros et al. 2010; Guo et al. 2020). *Spodoptera frugiperda* is considered the key pest of corn in Brazil and the spinosyn insecticides are one of the chemical groups used to manage this pest in this crop (Burtet et al. 2017). On the other hand, spinetoram is not registered to control *S. frugiperda* in soybean, and the inclusion of this insecticide for *S. frugiperda* control in cotton was approved in Brazil in 2020 (MAPA, 2021). Therefore, one hypothesizes is that the continuous selection pressure by spinetoram exerted on corn crops selects not only the resistant individuals to the insecticide molecule but also the most adapted individuals in this host plant. While the low or no use of spinetoram in soybean and cotton plants could explain the less adaptability of RR strain in these two host plants.

The slow development of *S. frugiperda* strains on soybean and cotton plants could also be related to the secondary compounds of these plants. The plant defense compounds and its nutritional quality affects the food conversion efficiency, which can increase the development time and fecundity of insect pests (Awmack and Leather 2002). Peruca et al. (2018) have demonstrated that secondary compounds of soybean plants have impaired the development of *S. frugiperda*. One of the most abundant secondary component of soybeans is the flavonoids, which were associated to behavior and physiological effects on insect pests (Piubelli et al. 2003, 2005; Bentivenha et al. 2018). In cotton plants, terpenoid, tannin and flavonoids have been reported as protective compounds against herbivorous insects and other organisms (Ti and Zhang 2009). But most of the studies in cotton are related to gossypol, a polyphenolic aldehyde from cotton that increases the fitness cost associated with Bt resistance in *Pectinophora gossypiella* (Carrière et al. 2004, 2019; Williams et al. 2011) and it might be one of the reasons for the lower fitness of RR strain in our study.

Herbivory defense compounds are also present in corn plants. Benzoxazinoids are defense compounds found in corn plants that have anti-digestive and toxic effects against insect pests (Wouters et al. 2016; Qi et al. 2018). DIMBOA is considered one the main benzoxazinoids in corn plants (Zhou et al. 2018) and several studies showed the effect of this compound on lepidopteran pests (Campos et al. 1989; Houseman et al. 1992; Ortego et al. 1998; Rostás 2007; Glauser et al. 2011). However, apparently DIMBOA did not affect the development of *S. frugiperda* (Rostás 2007; Glauser et al. 2011; Wouters et al. 2014). Rostás (2007) suggested that *S. frugiperda* appears to be better adapted to DIMBOA than *S. exigua* and use this secondary compound for host recognition and as a feeding stimulant. This controversy between a toxic secondary component of corn and the absence of fitness cost in *S. frugiperda* has been recently studied. Researchers showed the involvement of UDP-glycosyltransferases in detoxification of DIMBOA by *S. frugiperda* (Israni et al. 2020; Silva-Brandão et al. 2021) and allied it to the coevolutionary relationship between them (Mello and Silva-Filho 2002).

Insects have evolved different strategies to overcome insecticides and plant secondary compounds. Detoxification enzymes are the main mechanism of insect adaptation to synthetic and natural xenobiotics (Li et al. 2007a; Schuler 2011; Feyereisen 2012; Heidel-Fischer and Vogel 2015; Lu et al. 2020; Vandenhole et al. 2020). UDP-glycosyltransferases and cytochrome P450 monooxygenases play an important role in the detoxification of flavonoids and terpenoids in insects (Luque and O’Reilly 2002; Huang et al. 2008; Krempl et al. 2016; Jin et al. 2019). The strains of *S. frugiperda* tested in this study probably needed to overcome the plant defense through detoxification enzymes in order to survive and complete their life cycles in each host plant. In this way, the energy reallocation to produce such detoxification enzymes to metabolize the allelochemicals of soybean and cotton might be associated to the difference in the survival and development time of *S. frugiperda* strains in our study.

Therefore, another hypothesize is that the lower fitness of the RR strain on plants of soybean and cotton are due to an interaction of the resistance mechanisms of spinosyn resistance in *S. frugiperda* and the detoxification mechanism to the plant defenses compounds of soybean and cotton. It is worth mentioned that this hypothesis does not exclude the other proposition discuss above. The lower fitness of RR strain in soybean and cotton can be related both to an extrinsic factor as insecticide extensive use in the crop and/or an intrinsic factor such *S. frugiperda* physiology. The resistance mechanisms to spinosyns have been studied for several species of insect pests. Target site insensitivity has been associated with spinosyn resistance in most cases (Perry et al. 2007; Baxter et al. 2010; Hsu et al. 2012; Silva et al. 2016; Zimmer et al. 2016; Wan et al. 2018; Wang et al. 2020a, b; Zuo et al. 2020). However, some studies showed that metabolic detoxification enzymes also could be involved in spinosyn resistance (Wang et al. 2009; Sial et al. 2011; Rehan and Freed 2014). At the moment, the mechanism underlying spinosyn resistance in *S. frugiperda* is still unknown. Multi-omics approaches will be considered in our future studies in order to identify the molecular mechanism associated with spinosyn resistance in *S. frugiperda*.

In conclusion, we showed that the rate of spinetoram resistance evolution in *S. frugiperda* might be dependent on the host plant. When feeding on plants of corn, the resistance may evolve more rapidly than when feeding on soybean and cotton plants. The shorter development time on corn plants increases the number of generations of the pest and consequently increasing the frequency of resistance alleles in the field. In plants of soybean and cotton the frequency of resistant individuals tend to be lower due to the presence of fitness cost. Despite the competitive disadvantage of resistant individuals on plants of soybean and cotton, an attention should be given to the heterozygous individuals because its performance is similar to the susceptible individuals regardless the host plant. The heterozygous individuals are the main carriers of resistance alleles and its competitiveness guarantee the permanence of resistance alleles in the field (Roush and Daly 1990). The information provided here support the design of effective IRM strategies, highlighting the importance to consider not only the biological aspects of the pest but also the host plants that are part of the agricultural system where the pests are found.

## Acknowledgments

We thank São Paulo Research Foundation (FAPESP) for granting a PhD scholarship to the first author (grant #2019/06217-8).

## Declarations

### Author Contribution Statement

RHK, ASG and CO conceived and designed the experiments. RHK, ASG, FEOP and JGR performed the experiments and collected the data. RHK conducted the statistical analysis. All authors interpreted and discussed the results. RHK wrote the manuscript with inputs from all authors; CO supervised the research.

### Funding

This work was supported by São Paulo Research Foundation (FAPESP) (grant #2019/06217-8).

### Conflict of Interest

The authors declare that they have no conflict of interest.

### Compliance with ethical standards

“This article does not contain any studies with human participants or animals performed by any of the authors.”

## Notes

### Competing Interest Statement

The authors have declared no competing interest.

